# Exploring the Druggable Conformational Space of Protein Kinases Using AI-Generated Structures

**DOI:** 10.1101/2023.08.31.555779

**Authors:** Noah B. Herrington, David Stein, Yan Chak Li, Gaurav Pandey, Avner Schlessinger

## Abstract

Protein kinase function and interactions with drugs are controlled in part by the movement of the DFG and ɑC-Helix motifs, which enable kinases to adopt various conformational states. Small molecule ligands elicit therapeutic effects with distinct selectivity profiles and residence times that often depend on the kinase conformation(s) they bind. However, the limited availability of experimentally determined structural data for kinases in inactive states restricts drug discovery efforts for this major protein family. Modern AI-based structural modeling methods hold potential for exploring the previously experimentally uncharted druggable conformational space for kinases. Here, we first evaluated the currently explored conformational space of kinases in the PDB and models generated by AlphaFold2 (AF2) (1) and ESMFold (2), two prominent AI-based structure prediction methods. We then investigated AF2’s ability to predict kinase structures in different conformations at various multiple sequence alignment (MSA) depths, based on this parameter’s ability to explore conformational diversity. Our results showed a bias within the PDB and predicted structural models generated by AF2 and ESMFold toward structures of kinases in the active state over alternative conformations, particularly those conformations controlled by the DFG motif. Finally, we demonstrate that predicting kinase structures using AF2 at lower MSA depths allows the exploration of the space of these alternative conformations, including identifying previously unobserved conformations for 398 kinases. The results of our analysis of structural modeling by AF2 create a new avenue for the pursuit of new therapeutic agents against a notoriously difficult-to-target family of proteins.

**Significance Statement:** Greater abundance of kinase structural data in inactive conformations, currently lacking in structural databases, would improve our understanding of how protein kinases function and expand drug discovery and development for this family of therapeutic targets. Modern approaches utilizing artificial intelligence and machine learning have potential for efficiently capturing novel protein conformations. We provide evidence for a bias within AlphaFold2 and ESMFold to predict structures of kinases in their active states, similar to their overrepresentation in the PDB. We show that lowering the AlphaFold2 algorithm’s multiple sequence alignment depth can help explore kinase conformational space more broadly. It can also enable the prediction of hundreds of kinase structures in novel conformations, many of whose models are likely viable for drug discovery.

## Main Text Introduction

Protein kinases are key regulators of numerous cell-signaling pathways via phosphorylation of substrates and are involved in a variety of processes, such as cell proliferation, movement, and growth, as well as immunological responses (3-5). Protein kinases have been prominent targets for modern drug discovery against a variety of diseases (6-8). As of January 2023, 72 inhibitors were FDA-approved that primarily target two dozen different kinases (7).

However, kinases are difficult to target selectively (9, 10). The ATP-binding site, which most inhibitors bind, is highly conserved (11, 12), and druggable allosteric sites can be difficult to identify and target. Additionally, kinase structures are highly flexible, adopting different pharmacologically relevant states (13, 14), particularly in the ATP-binding site, which includes the DFG (Asp-Phe-Gly) motif. The ‘DFG-in’ conformation facilitates ATP binding (15), whereas the ‘DFG-out’ state, characterized by a roughly 180° ‘flip’ of the Asp and Phe residues, precludes ATP binding (13, 14, 16). The conformational plasticity of kinases is also characterized by the movement of the nearby ɑC-Helix, which can also adopt ‘in’ and ‘out’ states (17).

The conformations of kinases are of the utmost importance for pharmacology and drug discovery. Kinase inhibitors, including multiple prescription drugs, have been classified based on the conformation to which they bind, and different inhibitor *types* show different biochemical and pharmacological profiles (18). For instance, Type-II inhibitors targeting the ɑC-Helix-in / DFG-out (CIDO) conformation, an inactive conformation, exhibit longer residence times compared to Type-I inhibitors, which target the active ɑC-Helix-in / DFG-in (CIDI) conformation (19, 20).

Furthermore, both DFG-out and ɑC-Helix-out conformations have additional, nearby allosteric pockets that can be targeted for drug discovery (21-25). Recently, additional conformations have been described, which represent intermediate states that may enable the development of a unique class of kinase inhibitors (26, 27). The design of small molecule kinase inhibitors is aided by structural data of kinases in specific conformations they are intended to bind (28).

A major limitation in the development of conformation-specific kinase inhibitors (e.g., Type-II) is the relatively small number of inactive structures in the PDB (29) available for rational drug design (27, 30). However, computational modeling has helped researchers bridge the gap in our structural knowledge of kinase conformations for drug design. For example, homology modeling-(31) and molecular dynamics (MD) simulations-(32) based methods have been applied to visualize alternative kinase conformations and advance the design of compounds targeting them (33, 34). However, homology modeling does not model long regions that are unaligned to template structures (35). Similarly, MD simulations often fail to sample the pharmacologically relevant ‘DFG flip’ due to time and forcefield limitations, which limit their utility in kinase drug discovery.

Emerging *ab initio* modeling methods address some of these barriers by employing machine learning (ML) algorithms to generate 3D models with accuracies comparable to experimentally determined structures (1, 2, 36, 37). Some of these methods, including AlphaFold2 (AF2) (1) and RosettaFold (36), use a multiple sequence alignment (MSA) as input that captures conserved contacts between evolutionarily related sequences needed to preserve the overall fold and topology governing all proteins (38). Another example is ESMFold (2), which is an alignment-free, language-based model capable of predicting protein structures based on biological properties learned directly from sequence data.

Recent work has demonstrated that the modulation of some of the AF2 prediction parameters may help explore structural variations in membrane proteins (39) and multidomain electron transfer complexes (40). In this work, motivated by the importance of protein kinases and their unique conformations, we examined the ability of two representative AI-based structural modeling methods, i.e., AF2 and ESMFold, to explore the kinase conformational landscape relevant to drug discovery. Focusing on AF2, we also evaluated the quality of predicted models with respect to known experimental data and devised an approach to promote conformational exploration of the DFG and ɑC-Helix motifs within these models. Our analyses suggest that a number of high-quality AF2 models that we predicted for several kinases were in never-before-seen conformations that can be used for drug design in future work.

## Results

### PDB Structures and AlphaFold2 and ESMFold Models Overrepresent the Kinase Active State

We first sought to assess the current conformational space of experimentally determined structures of kinases in the PDB and structural models deposited in the AlphaFold2 Protein Structure Database (‘AF2 Database’) (41) and generated by ESMFold (2). We downloaded all available experimentally determined structures of 497 human kinase domains (from 484 kinases) from the PDB. Although, on average, each kinase has more than 10 structures in the PDB, many kinases (268) have no known experimentally determined structure. We found that, as of May 9^th^, 2023, the PDB had 5,136 structures classifiable (42) into one of six defined conformations, including ɑC-Helix-in / DFG-in (CIDI), ɑC-Helix-in / DFG-out (CIDO), ɑC-Helix-out / DFG-in (CODI), ɑC-Helix-out / DFG-out (CODO), DFGinter, and Unassigned (Figure 1A). These structures covered 270 kinases, of which 78.9% only exhibited two or fewer conformations. A summary of our workflow is visible in Supp. Figure 1.

**Figure 1.**
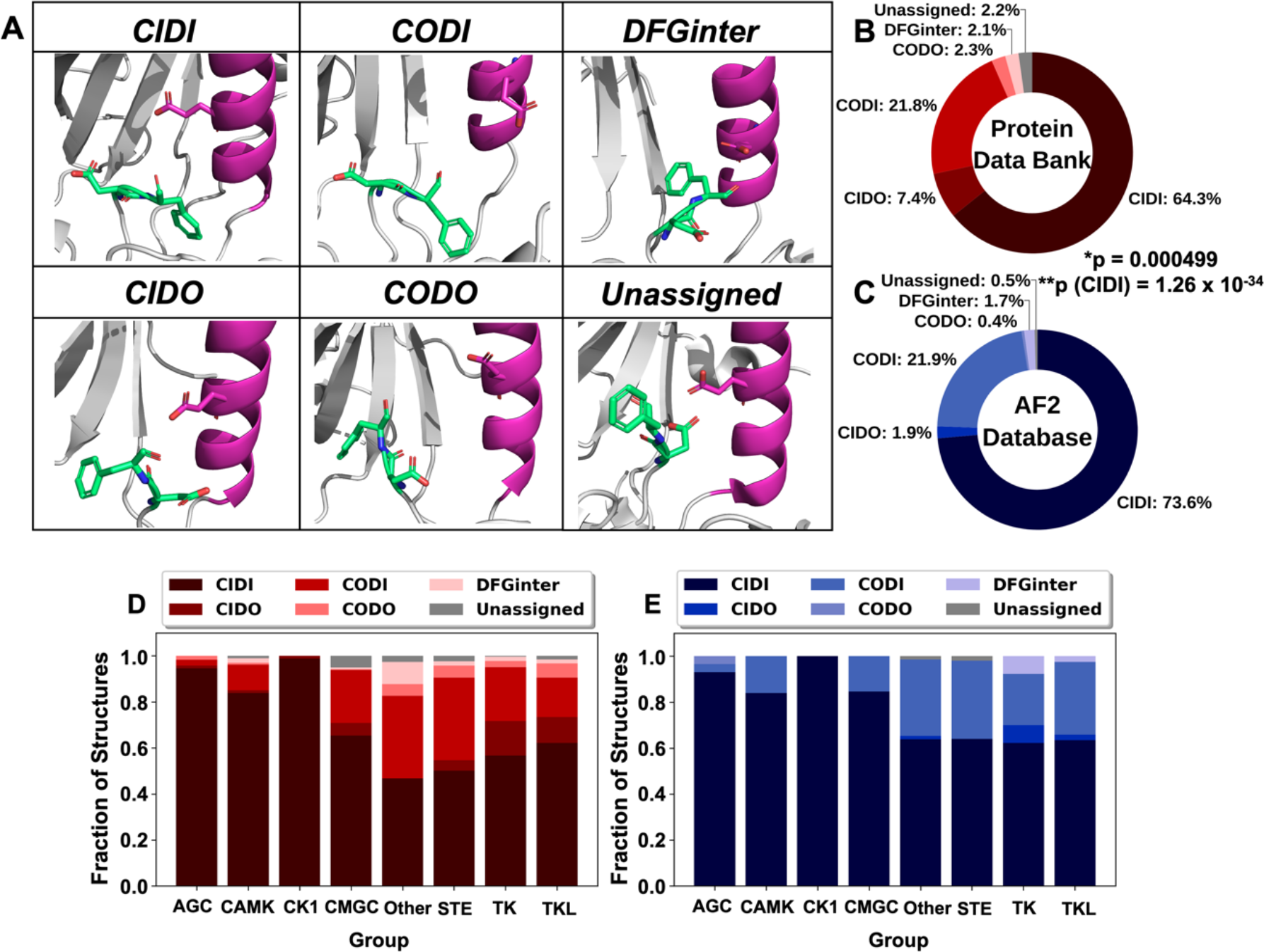
Kinase conformations in the PDB and AlphaFold2 Predicted Protein Structure Database. **A)** Prototypical structural conformations considered in our study: CIDI: ɑC-Helix/DFG-in; CIDO: ɑC-Helix-in/DFG-out; CODI: ɑC-Helix-out/DFG-in; CODO: ɑC-Helix-out/DFG-out; DFGinter; Unassigned Examples, with PDB IDs, shown are CIDI: MAPK14 (1BL6), CIDO: RIPK2 (4C8B), CODI: CDK2 (1H07), CODO: NTRK1 (6D1Y), DFGinter: AURKA (3FDN), Unassigned: PDGFRA (6JOJ). Residues in lime green indicate the aspartic acid and phenylalanine (DF) of the DFG motif, while magenta indicates the ɑC-Helix, whose conformation is signaled by movement of the conserved Glu residue. Red and blue atoms indicate oxygen and nitrogen atoms, respectively. **B)** Fractional representation of kinase structures from the PDB (n=5,024) classified into each conformation type. The plot indicates an over-representation of the CIDI state compared to other conformations. **C)** Fractional representation of kinase domain models from the AF2 Database (n=469) classified into each conformation type. *p-value was calculated using the Fisher’s exact test and indicated that the PDB and AF2 distributions were different. **p(CIDI) was calculated using a one-sided Wilcoxon rank-sum and indicated that AF2 had a higher overrepresentation of CIDI models compared to the PDB. **D)** Fractional distributions of models in different conformations by kinase group (AGC: PKA, PKG, PKC families; CAMK: Calcium/calmodulin-dependent; CK1: Casein kinase 1; CMGC: CDK, MAPK, GSK3, CLK families; STE: Sterile 7, Sterile 11, Sterile 20 kinases; TK: Tyrosine kinase; Tyrosine kinase-like) in the PDB. CIDI models dominated the distribution for each group, while DFG-out conformations (CIDO and CODO) are significantly underrepresented. Unassigned models tended to be rare. **E)** Fractional distributions of models in different conformations by kinase group in the AF2 Database. CIDI models dominated across all groups here as well, while DFG-out models were much less common and even rarer than in the PDB. Unassigned models were also rarer than in the PDB.

We also downloaded all computational models of these domains from the AF2 Database and predicted structures of the same kinase domains using ESMFold, which were all also classified by their conformation. Included in the AF2 Database were models of 153 kinases with no deposited structures in the PDB. We calculated the fractional distribution of the six conformation types present in the PDB and AF2 Database (Figures 1B and 1C) and models predicted by ESMFold (Supp. Figure 2). Our analysis revealed a significant overrepresentation of the active state ‘CIDI’ in all three datasets. Interestingly, both AF2 and ESMFold exhibited an even greater fraction of active-state models than that of the PDB (73.6%, 82.9% and 65.7%, respectively; p(PDB_CIDI_ < AF2_CIDI_) = 1.26 x 10^-34^; p(PDB_CIDI_ < ESMFold_CIDI_) < 2.2 x 10^-16^. Furthermore, DFG-out conformations were underrepresented in all three datasets, more so in the AF2 Database and by ESMFold than in the PDB (2.3%, 0.8% and 9.7%, respectively; p(PDB_DFG-out_ > AF2_DFG-out_) < 2.2 x 10^-16^; p(PDB_DFG-out_ > ESMFold_DFG-out_) < 2.2 x 10^-16^). Notably, all three datasets contained a small fraction of Unassigned models (PDB: 2.2%; AF2: 0.4%; ESMFold: 0.4%). After limiting the analysis to models of kinases with deposited structures in the PDB, the distributions of AF2 and ESMFold model conformations of kinases showed similar trends when compared to those of their full datasets (Supp. Figure 3; p(AF2 Overlap) = 0.751; p(ESMFold Overlap) = 0.913), indicating that these differences do not arise from the expanded coverage of the kinome by AF2 or ESMFold. This suggests that these *ab initio* predictors exhibit a bias for predicting structures of kinases in the active (CIDI) state, which should be addressed to expand the conformational exploration of the kinome.

**Figure 2.**
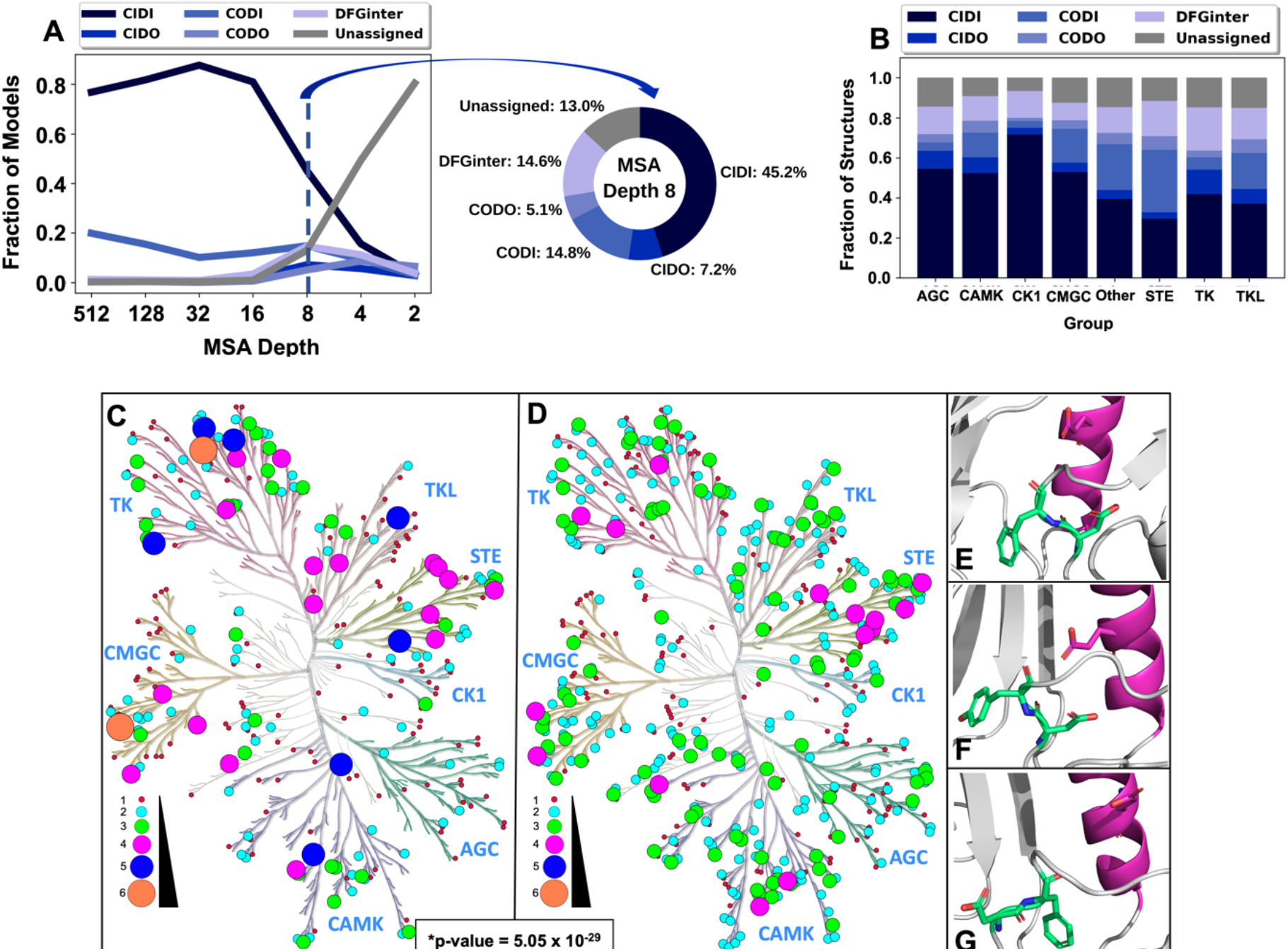
Exploration of the kinome’s conformational space by lowering MSA depth when predicting structures using AlphaFold 2 (AF2). **A)** Fractional distribution of models in various conformations across different MSA depths (left plot) and at an MSA depth of 8 (pie chart). AF2 models were variably distributed across the various conformations at different MSA depths, the highest of which yielded large fractions of CIDI models among their predictions. Lower MSA depths, primarily beginning at a depth of 8, yielded higher fractions of non-CIDI (inactive state) models. **B)** Distributions of models predicted at an MSA depth of 8 by kinase group. Certain groups, including TK and TKL, exhibited relatively high fractions of DFG-out (CIDO and CODO) models (16% and 14%, respectively) compared to those predicted by AF2 under default parameters. Unassigned model fractions were also greater among all groups when compared to the full set of AF2 structures (Supp. Table 3). **C)** Coverage and conformational space representation across the human kinome tree, generated with KinMap, for structures from the PDB, where each node represents one kinase. Colors (crimson, cyan, lime, magenta, blue and coral, in order) and increasing circle sizes indicate greater counts of unique conformations. More than 88% of kinases in the PDB have been solved in only two or fewer unique conformations. **D)** Coverage and conformational space representation of the human kinome using models predicted by AF2 at an MSA depth of 8. The node color and size coding is the same as in **C**. 46% of the models exhibited two or fewer unique conformations at this MSA depth. In comparing paired counts of kinase conformations between both datasets, on average, each kinase exhibits more unique conformations at this depth than in the PDB (p-value = 5.05 x 10^-29^). **E-G)** Representative models predicted by AF2 at an MSA depth of 8 for kinases from different groups in novel conformations not seen in the PDB. Examples shown are **E)** CLK3 in the CIDO conformation, **F)** NEK9 in the CIDO conformation, and **G)** MAP3K4 in the CODI conformation. Residues in lime green represent the ‘DF’ of the DFG motif (‘DY’ of DYG for NEK9), while residues in magenta represent the ɑC-Helix. Red and blue atoms indicate oxygen and nitrogen atoms, respectively.

**Figure 3.**
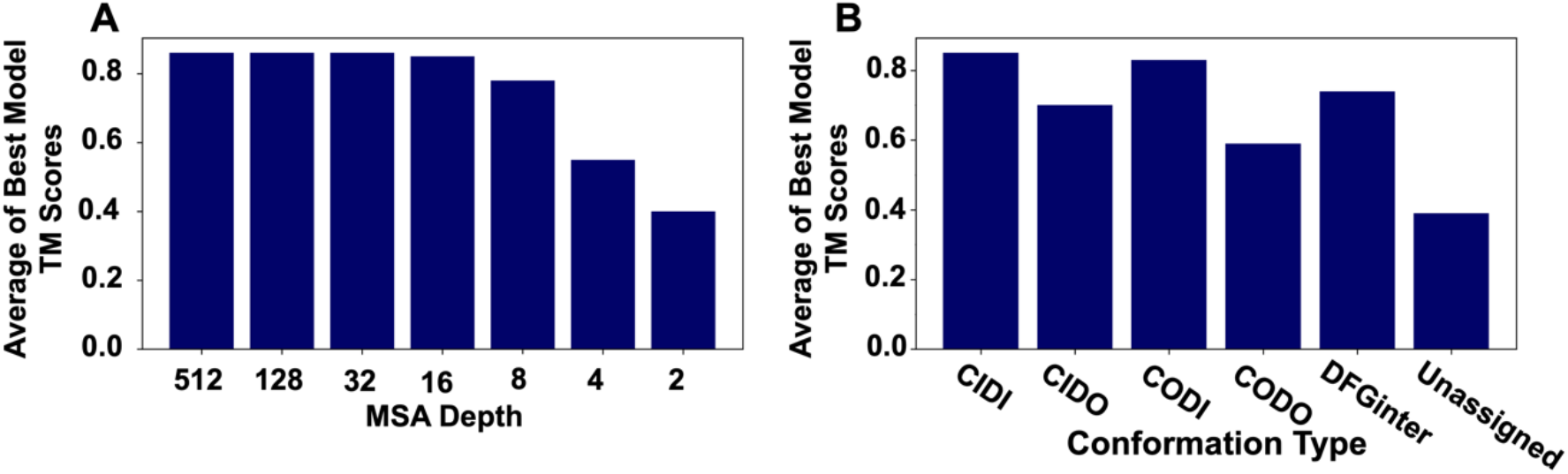
Accuracy of AF2 models by MSA and conformation type. Bar plots summarizing the accuracy in terms of TM-Scores calculated for AF2 models by comparing them to the corresponding PDB structures. **A)** Averages of the highest TM-Scores calculated from comparisons between all AF2 models to all PDB structures of the same kinase by MSA depth. A trend of decreasing model accuracy is evident, especially from an MSA depth of 16 through 2. **B)** Averages of the best TM-Scores calculated from comparisons between all AF2 models to all PDB structures of the same kinase by conformation type. CIDI models were the most accurate, followed by those in CODI and DFGinter conformations. CIDO and CODO models were generally less accurate than DFG-in models. Unassigned models were the least accurate.

### Different Kinase Groups Tend to Adopt Distinct Conformations

Next, we investigated if the bias for the CIDI conformation existed within individual kinase groups. We therefore computed the fractional distributions of each conformation by kinase group (AGC, CAMK, CK1, CMGC, Other, RGC, STE, TK and TKL) (43) using our datasets from the PDB and AF2 Database (Figures 1B and 1C) and the models generated by ESMFold. We observed that the active conformation, CIDI, consistently represented the greatest fraction among all kinase groups in all three datasets, ranging from 46.7% (Other) to 98.9% (CK1) in the PDB, 62.2% (TK) to 100% (CK1) in the AF2 Database, and 58% (STE) to 100% (CK1 and CMGC) by ESMFold (Supp. Figure 4). Interestingly, the CAMK and CK1 groups appeared to have similar fractions of each conformation type between the PDB and AF2 Databases (CAMK: p = 0.587; CK1: p = 1). These same groups did not retain those similar distributions in ESMFold-predicted models (Supp. Figure 4).

**Figure 4.**
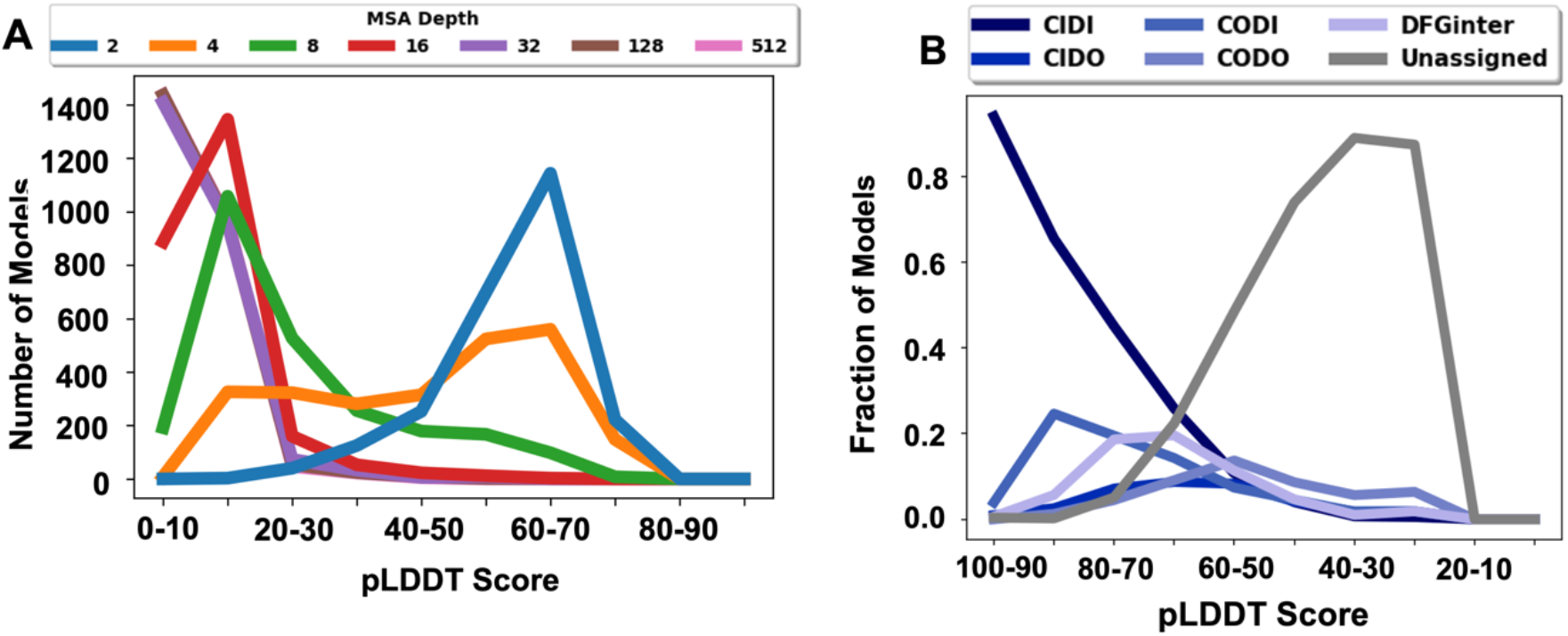
Distributions of accuracies of AF2-generated models in terms of the pLDDT score. pLDDT, as output by AF2, is a measure of confidence in the predicted model, calculated as a whole-chain average of per-residue scores of predicted model accuracy. **A)** Distribution of kinase models by pLDDT across different MSA depths. High MSA depths typically yielded more confident models, as low MSA depths typically yield less confident models. **B)** Distribution of pLDDT scores of all models predicted across all MSA depths, grouped by conformation type. CIDI models represented the most confident models. CIDO, CODI, CODO and DFGinter models more broadly spanned different levels of confidence, while Unassigned models accounted for the vast majority of low-confidence predicted structures.

Other groups showed noticeable differences. For example, those groups having higher fractions of CIDO structures in the PDB, such as CMGC, STE, TK, and TKL, had lower CIDO fractions by AF2 and ESMFold. However, it is noteworthy that TK had the highest CIDO fraction in all three datasets, since previous analyses have shown members of this group to be more likely to adopt DFG-out states (15, 43). Intriguingly, the understudied conformation CODO was a minor population across all groups in the PDB, ranging from 0.0% (CK1) to 6.1% (TKL), but was even less frequent in the AF2 and ESMFold datasets, ranging from 0.0% (all but AGC) to 3.4% (AGC) by AF2 and 0.0% (all but AGC) to 1.7% (AGC) by ESMFold. Finally, in addition to the sole CODO model belonging to AGC generated by ESMFold, the only two Unassigned models belonged to the Other group, and the only three CIDO models belonged to TK. Taken together, our results suggest that the conformation bias seen across the kinome (Figure 1B) is also observed at the group level for the PDB and both predictive methods and that some groups are more conformationally diverse than others.

### Lowering MSA Depth Increases the Structural Diversity of AF2-Generated Kinase Models

The AF2 algorithm includes a variety of parameters whose tuning can impact the output protein structural models. Specifically, the number of sequences (‘depth’) in the multiple sequence alignment (MSA) used as input for AF2 correlates with the conformational diversity of the structural models generated of a given sequence (39). Therefore, to sample alternative kinase conformations, we used ColabFold (44) to run AF2 at various MSA depths as input (Methods). The MSA generated by AF2 includes evolutionarily related sequences to the input kinase sequence. We generated five models for each kinase, which were then classified into the same six conformations as earlier. We observed limited structural diversity in models generated at higher MSA depths of 512, 128, and 32, where most distributions appeared similar to those observed in the AF2 dataset, though only the distribution at a depth of 512 was statistically similar (Figure 2A; Supp. Table 1). For example, at an MSA depth of 512, 76.8% of the models were in the active conformation, CIDI, while DFG-out states (CIDO and CODO) were under-represented (total of 1.7%).

**Table 1.** Counts of confidently predicted AF2 models in conformations (columns) not seen for the same kinases in the PDB at multiple MSA depths (rows).

**pLDDT > 70 (pLDDT > 90)<sup>b</sup>**
| **MSA<sup>c</sup>** | **Kinases<sup>d</sup>** | **All<sup>e</sup>** | **CIDI** | **CIDO** | **CODI** | **CODO** | **DFGinter** | **Unassigned** |
| --- | --- | --- | --- | --- | --- | --- | --- | --- |
| **2** | **25 (0)** | **30 (0)** | **6 (0)** | **3 (0)** | **7 (0)** | **2 (0)** | **11 (0)** | **1 (0)** |
| **4** | **275 (2)** | **433 (2)** | **119 (0)** | **33 (0)** | **88 (0)** | **44 (1)** | **122 (1)** | **27 (0)** |
| **8** | **354 (68)** | **1066 (94)** | **480 (90)** | **98 (1)** | **180 (3)** | **49 (0)** | **236 (0)** | **23 (0)** |
| **16** | **278 (127)** | **1220 (406)** | **933 (395)** | **31 (0)** | **169 (11)** | **11 (0)** | **67 (0)** | **9 (0)** |
| **32** | **244 (153)** | **1197 (645)** | **1008 (620)** | **20 (4)** | **147 (16)** | **0 (0)** | **14 (1)** | **8 (4)** |
| **128** | **239 (160)** | **1178 (653)** | **918 (589)** | **17 (17)** | **215 (29)** | **1 (0)** | **18 (9)** | **9 (0)** |
| **512** | **239 (153)** | **1173 (645)** | **838 (557)** | **22 (21)** | **285 (48)** | **4 (0)** | **18 (13)** | **6 (6)** |

<sup>a</sup> 'Unique Kinases All Depths' signifies the total count of all sequence-unique kinases predicted in novel conformations across all MSA depths.
<sup>b</sup> Numbers given in parentheses represent the same counts for models with pLDDT scores higher than 90 to assess the robustness of the findings for even higher accuracy models.
<sup>c</sup> 'MSA' indicates the MSA depth or the number of sequences as input to AF2 to predict structural models of kinases.
<sup>d</sup> 'Kinases' corresponds to the number of unique kinases predicted in novel conformations at each MSA depth.
<sup>e</sup> 'All' corresponds to the total count of models predicted in novel conformations for their respective kinases at each MSA depth.

Interestingly, we observed greater fractions of alternative conformations in models generated at shallow MSAs (i.e., MSAs with fewer sequences) (Figure 2A). For instance, at an MSA depth of 8, the CIDI fraction of models was 45.2%, while the DFG-out states’ (CIDO and CODO) fraction was 12.3%. At lower MSA depths of 4 and 2, the models were even more diverse, respectively including only 15.7% and 3.2% of the models in the CIDI conformation. Furthermore, substantial fractions of the models (49.3% and 80.5%, respectively) were classified as Unassigned at these MSA depths. Statistical comparison of the fractional compositions of each conformation at each MSA depth to each other showed that they were all statistically different (Supp. Table 2). We also examined if the conformational predisposition of some sequences can affect other model predictions. For this, we used custom MSAs made only from sequences predicted to be in at least one DFG-out state out of five models at an MSA depth of 8. These custom MSAs were generated from all such 161 kinases, with depths ranging from 161 to 4 sequences, to favor prediction of structures in those inactive conformations. However, these models exhibited similar conformational distributions to that observed in the AF2 Database (Supp. Figure 5), i.e., favoring the CODI and CIDI DFG-in conformations (Methods).

**Figure 5.**
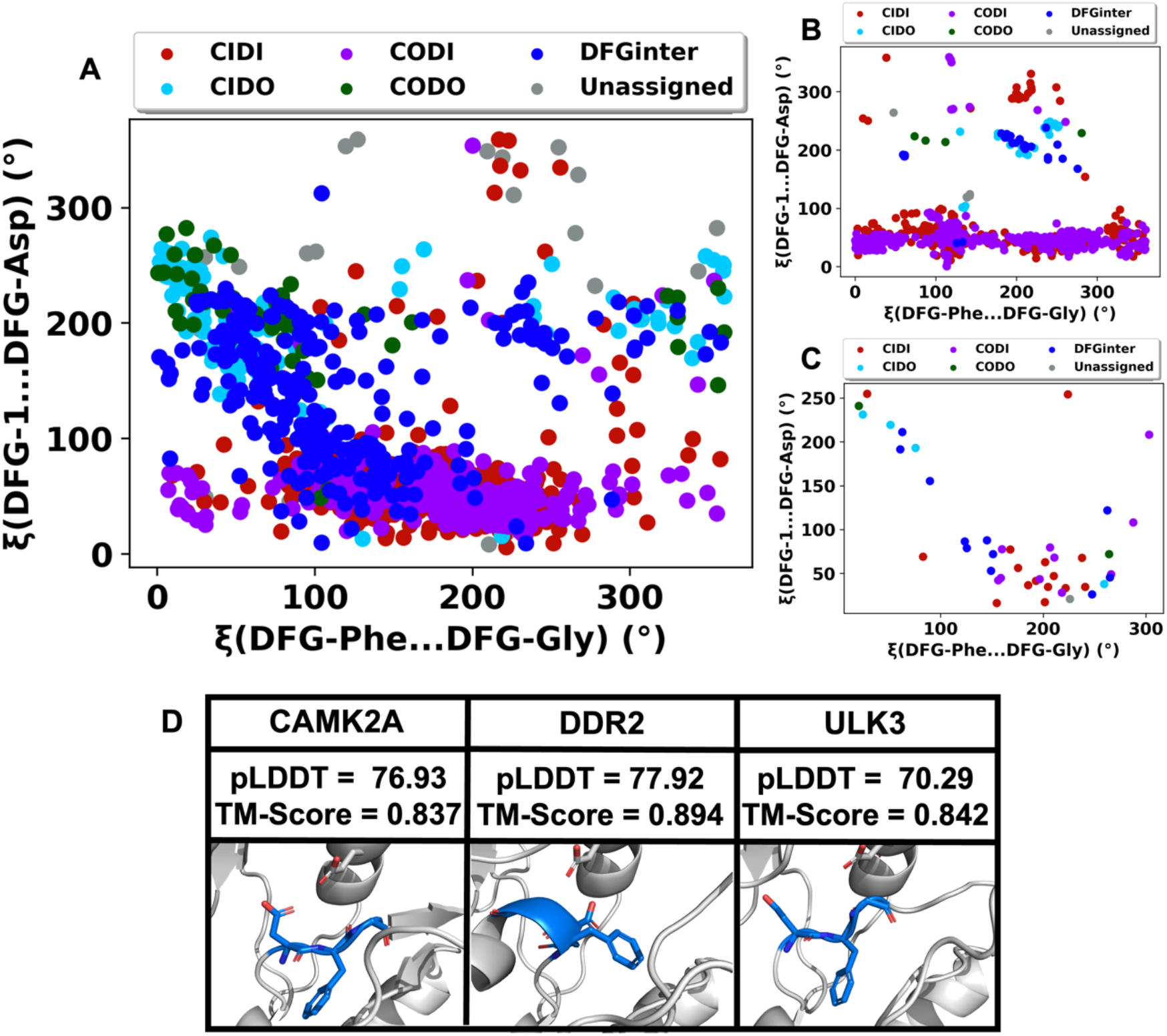
Distribution of AF2-predicted models in terms of the movement of the DFG motif, as defined by pseudo-dihedral angles, for AF2-predicted models. For models predicted at MSA depths of (A) 8, (B) 512, and (C) 2, pseudo-dihedral angles are defined by Möbitz (49). The X-axis is defined by the pseudo-dihedral angle constructed by the Cɑ carbons of the DFG-Asp, DFG-Phe, DFG-Gly and DFG+1 residues, while the Y-axis is defined by the angles of the DFG-2, DFG-1, DFG-Asp and DFG-Phe residues. **A)** Distribution of all models predicted at an MSA depth of 8 with pLDDT > 70, categorized by their adopted conformation. **B)** As defined in A, dihedral angles plot of all AF2 models with pLDDT > 70 predicted at an MSA depth of 512. Movement of the DFG motif into DFGinter and DFG-out conformations and overall conformational exploration were much more limited at this depth than at 8. **C)** Dihedral angles plot of all AF2 models predicted at an MSA depth of 2 with pLDDT > 70. Movement of the DFG motif into diverse conformations was much more frequent at this MSA depth, but far fewer models were confidently predicted. The majority of confident models were still DFG-in at this depth. **D)** Examples of confident ‘Unassigned’ models are illustrated, where red and blue atoms indicate oxygen and nitrogen atoms, respectively.

Additionally, we observe that lowering MSA depth impacts the distribution of model conformations by kinase group (Figure 2B). First, the CIDI fraction at an MSA depth of 8 is much lower for all groups than the corresponding fractions in the AF2 Database, and ranges from 29.6% (STE) to 71.7% (CK1) (Figures 1E and 2B). Second, the fractions of CIDO models in most groups is higher at an MSA depth of 8 (3.3% (CK1) to 12.1% (TK)) than the corresponding fractions in the AF2 Database (0.0% (many) to 7.8% (TK)). A similar trend is observed in CODO models for all groups (range of 1.7% (CK1) to 6.8% (STE) at MSA depth of 8), except AGC. The levels of statistical significance of all these comparisons are provided in Supp. Table 3.

To better understand these findings, we visualized the number of unique conformations for each kinase using KinMap (45), a tool for presenting data across the kinome, for the PDB structures and AF2 (MSA depth=8) models (Figures 2C and 2D, respectively). A comparison between these kinome trees demonstrated that lowering the MSA depth explored the conformational space of the kinome more broadly than the PDB and that, on average, an MSA depth of 8 yielded more conformations predicted per kinase over those with structures in the PDB. (p-value = 5.05 x 10^-29^). Part of this expansion is in the form of conformations not previously reported for many kinases. For instance, CLK3, NEK9, and MAP3K4, all from different kinase groups, have no experimentally determined structures in the PDB but were predicted confidently in previously uncharacterized conformations (CIDI, CIDO, and CODI (CLK3); CIDI and CIDO (NEK9); CIDI, CODI, CODO, and DFGinter (MAP3K4)) by AF2. Collectively, the above data suggest that conformational exploration is not limited to any one kinase group.

### Models Generated by AF2 at Different MSAs Vary in Quality

While the above results demonstrated that AF2 can generate previously unseen kinase conformations, an assessment of model quality is a prerequisite for their utility for drug discovery and other applications. To evaluate the models’ quality, we compared them to experimentally determined structures from the PDB using the template modeling-score (TM-Score) (46), which is commonly used to evaluate protein structure prediction methods (1, 47) (Methods). We focused on models specifically containing the DFG motif. Briefly, we aligned each predicted model with every structure of the same kinase in the PDB and calculated a TM-Score of the aligned structures. To consider the comparison of each model to the most closely matching PDB structure, this was followed by calculating an average of the best TM-Scores for all models at each MSA depth and for each conformation type (Figure 3). We observed that the average TM-Score for models deteriorated with MSA depth starting at a depth of 16 (0.85 to 0.4 at MSA depth 2) (Figure 3A). These results indicated that while lower MSA depths did explore a larger conformational space, they were also likely to lead to models of lower accuracy.

In terms of individual structural conformations, we observed that DFG-in models exhibited better accuracy scores than those in the DFG-out and DFGinter conformations (Figure 3B). For instance, CIDI models were the most similar to known structures (TM-Score of 0.85), while Unassigned models were the least (TM-score of 0.39). Notably, models classified as DFG-out (CIDO and CODO) tended to be more dissimilar to PDB structures (TM-scores of 0.7 and 0.59, respectively) than CIDI models.

One limitation of comparing models to PDB structures using similarity metrics like TM-score is that they penalize previously unobserved conformations that did not appear in the PDB. Therefore, we also calculated and analyzed the predicted local-distance difference test (pLDDT) scores (1, 41) for the AF2-generated kinase models. The pLDDT score assesses the overall confidence in AF2’s prediction on a per-residue level, ranging from 0 (low confidence) to 100 (high confidence), and is more lenient toward structural novelty than direct similarity metrics, such as TM-Score. We analyzed the distribution of average pLDDT scores taken from all residues in each model by MSA depth and conformation (Figure 4). Our analysis revealed that, as the MSA depth decreased from 512 to 2, we observed a nearly consistent drop in the overall confidence of models (Figure 4A).

We also observed that different conformations had varying levels of confidence in terms of the pLDDT score (Figure 4B). Models in the active CIDI conformation were predicted with the highest level of confidence (94% at 90 < pLDDT < 100), while Unassigned models were mostly predicted at very low levels of confidence (89% at 30 < pLDDT < 40). Other conformations were more broadly distributed across the pLDDT range.

Collectively these data suggest that, while AF2 did produce models of varying accuracies in different conformations, excessive lowering of the MSA depth diminished the overall quality of predicted models.

### AlphaFold2 Predicts Structures in Diverse DFG-in, DFG-out and Unseen Conformations

While we understood that high model quality was an important requirement for rational drug design using these models, it was also important to assess how well AF2 explored the conformational flexibility of the DFG motif. To do this, we next sought to evaluate the range of DFG motif movement captured by AF2. This movement determines the biophysical features (e.g., shape and volume) of the ATP binding site, which are relevant for the development of particular kinase inhibitor types (e.g., Type-I and -I½) (24, 25, 48). For this, we analyzed the reliable models generated (i.e., those models with pLDDT > 70) by AF2 at an MSA depth of 8 (Figure 5A) in terms of pseudo-dihedral angles involving the DFG backbone (49). Briefly, these pseudo-dihedral angles capture movement of the DFG motif and distinguish between DFG-in and DFG-out models. In Figures 5A the DFG-in conformations (red and purple) clustered toward the bottom of the plots where DFG-in models typically do, and DFG-out models (cyan and green) clustered in a line near the middle in the same way. DFGinter models (blue) spanned the conformational space between the two states. We observed that reliable Unassigned models, which were relatively uncommon at depth 8, often fell outside the regions occupied by the distinct DFG-in, DFG-out and DFGinter groups.

Notably, for the MSA depth of 512, we observed a limited exploration of DFG movement in the DFG-out and Unassigned conformational space (Figure 5B), indicated by narrower clusters of models in each region and fewer DFG-out models. In contrast, the MSA depth of 2 appears to show wider exploration of the DFG motif’s conformational space, but it is difficult to make definitive conclusions, as far fewer models were confidently predicted at this depth (Figure 5C), These observations demonstrate that reliable AF2-generated models captured the known conformational space of the DFG motif, but also some diverse and potentially druggable conformations (i.e., via the ATP binding site) outside of standard groupings.

### AlphaFold2 Predicts Structures of Kinases in Previously Unseen Conformations

Encouraged by AF2’s ability to predict inactive conformations of kinases, we investigated whether any kinases in our dataset were predicted in previously unobserved conformations. For this, we compared all models output by AF2 to PDB structures of the same kinases and calculated the number of models predicted in conformations not yet experimentally observed for their respective kinases at each MSA depth. We also included in our analysis models of kinases for which no structures currently exist in the PDB. We only considered ‘confident’ models, as defined by a pLDDT score higher than 70 (Table 1) (1, 41). A total of 6,297 confident models were generated in previously unobserved conformations, covering 398 kinases and 37% of all models generated across all MSA depths. This attests to our ability to explore new conformational spaces for many established and emerging drug targets that can now be targeted by inhibitors of different types in an expansion of the druggable kinome (Figures 2C and 2D). For example, confident (pLDDT > 70) models of CAMK2A, DDR2 and ULK3 at an MSA depth of 8 in Unassigned conformations represent previously unobserved and potentially druggable kinase models (Figure 5D). These three kinases reside in different groups and demonstrate that our predictions provide new structural information, not just for one family of kinases, but across the kinome. However, it should be noted that several of these models in new conformations contain partially obstructed ATP sites and DFG pockets. The seemingly collapsed binding sites likely make rational drug design challenging and may require further modeling to dock against them. To enable further study of and rational drug design efforts against such targets, all confident AF2-generated models analyzed here have been made available at https://kinametrix.com/ under the ‘AlphaFold2 Kinase Structures’ tab with helpful scripts through our GitHub (https://github.com/schlessingerlab/af2_kinase_conformations/).

## Discussion

A major barrier for rational design of conformation-specific kinase drugs is the limited availability of structures in diverse conformations. While inhibitors targeting the kinases’ ATP binding site may suffer from issues such as promiscuity and low residence time, kinase inhibitors targeting inactive states (e.g., Type II and Type 1½) can offer several advantages, such as improved pharmacokinetic properties and chemical novelty (18-20, 50). However, the limited number of structures of kinases in inactive states (e.g., DFG-out or ɑC-Helix-out) in the PDB attests to the difficulty of capturing these elusive conformations by experimental methods (Figure 1). Therefore, refined computational prediction methods are necessary to bridge this gap in our structural knowledge for the purpose of advancing drug discovery. Here, we performed a comprehensive analysis of the kinase conformational space captured by experimentally derived and AI-predicted structures, with the following three key findings:

### The PDB, as well as Default AlphaFold2 and ESMFold Models, Exhibit Bias for Active Kinase Conformations

Automated classification of kinase conformations can be achieved with high accuracy (42, 51), enabling high-throughput conformational analysis for kinase structure datasets. We determined fractions of kinase conformations in the PDB and AF2 Database, as well as models predicted by ESMFold (Figure 1). Our analysis revealed a significant bias within the PDB for the active kinase state (Figure 1B), which is consistent with previous analyses using different approaches (27, 42). This bias was even more pronounced in the AF2 Database and among models predicted by ESMFold (Figure 1C). Furthermore, DFG-out states represented only 10%, 2%, and <1% of the kinases in the PDB, AF2 Database, and ESMFold models, respectively, and together were only a minor fraction of structural data for conformation-specific ligand design. The bias in the models generated by both the AI-based methods likely resulted from the inherent bias in the PDB, which was used for their development and/or evaluation (1, 2). Alternatively, it is plausible that the methods favor the most energetically stable active (CIDI) conformation (16, 24). Future methods can potentially be trained and evaluated on structures of kinases in specific conformations to make the methods more representative. Encouragingly, although the handful of AI-predicted models in inactive conformations does not fully address the gap in our knowledge of kinase conformations, even a small number of high-quality models in unique conformations may enable the identification of targeted, selective compounds.

### Multiple Sequence Alignment (MSA) Depth Influences the Diversity of AlphaFold2-Predicted Kinase Structures

We explored whether varying the MSA depth, a core parameter of the AlphaFold2 algorithm, generated kinase models in inactive states. Our analysis showed that lower MSA depths correlated well with increased conformational diversity and yielded some high-quality models in previously unobserved conformations (Table 1). For example, NEK9 and CLK3 were predicted with high confidence in the CIDO conformation and MAP3K4 in the CODI conformation, which have not been seen for these proteins before (Figures 2E-G respectively). This suggests that shallow MSAs fed as input to AF2 result may result in fewer structural constraints, thereby enabling the method to sample rare, but possible, conformational states. Further studies should examine what information embedded in sequence alignments drives enhanced sampling of alternative conformations, as multiple custom alignments of kinases predicted in DFG-out states (CIDO and CODO) did not enhance sampling of those conformations at various depths; rather, all of these distributions were similar to that of models predicted by AF2 under default parameters (Supp. Figure 5).

However, our efforts did not fully overcome AF2’s bias for predicting kinases in active states, which could be based on AF2’s training on the PDB (52). This may also support the hypothesis that some kinases are less capable of adopting certain conformations, such as DFG-out, as has been suggested (53). It also further supports the notion that new and current computational tools should be developed to address this barrier to novel structure prediction.

### Several Predicted Models, Including Those in an Unassigned Conformation, May Represent A Novel Druggable Space for Kinases

Particularly interesting were the AF2-predicted kinase models *without* an assigned conformation, as these showcase the method’s potential to probe the unexplored conformational space of this family of proteins. While several high-scoring models were found in diverse structural conformations, a key question remained: were these models sufficiently accurate for the development of future novel kinase therapeutics? To answer this question, we performed a comprehensive analysis of AF2-generated kinase models and their druggable conformations. Although a large fraction of our models were of insufficient quality to be used in drug discovery, AF2 was able to capture a few high scoring models in unique conformations (Figure 5). Many of these models could represent new starting points for drug design. A large kinase inhibitor profiling study proposed that hundreds of kinases may bind Type-II inhibitors in the DFG-out state, far more than have been experimentally determined (28); from this, we may learn that screening for drug candidates need only identify ligands that effectively stabilize these potentially elusive conformations. However, a closer examination of some of the models revealed partially occluded binding sites, which necessitates further analysis and model refinement. For example, MD simulations on predicted protein structural models may be key to opening these pockets, as has been suggested before for homology modeling protocols (54). Moreover, our analysis indicated a delicate balance between the exploration of the structural conformational space and the attainment of high model quality (Figure 4).

Taken together, our results suggest that a rigorous and intentional interrogation of AI-predicted models by a trained scientist remains an important step in kinase drug design. While AI-based methods can make the structural characterization of kinases more efficient, the translation of these models into drug discovery campaigns still likely requires manual inspection to verify the placement of binding site sidechains, openness of druggable pockets, and large conformational changes (55). Nevertheless, our work here marks the beginning of the path towards the development of novel kinase therapeutics based on AI-generated structural models. Also, as more kinase structures are solved in alternative conformations and novel ligands targeting these states are developed, newly trained machine learning (ML) algorithms will likely be more generalizable, potentially further driving drug discovery.

### Limitations

As mentioned above, many of the generated kinase models contain occluded or partially occluded binding sites, or connectivity errors. The majority of the inspected cases will likely require further careful analysis by a trained modeler and model refinement (e.g., using MD) in order to be useful for drug discovery campaigns. In addition, our study is focused on investigating how changes to the input MSA affect distributions of kinase model conformations. Using additional templates or allowing input of multiple random seeds, which will be explored in future studies, may also affect conformational diversity of generated models.

## Materials and Methods

### AlphaFold2 Model Generation

High-throughput <u>AlphaFold2</u>model generation was performed using the Minerva high-performance computing facilities at the Icahn School of Medicine at Mount Sinai. An NVIDIAA100_PCIE_40GB GPU was used to run <u>the method</u>. Input sequences were derived from a previously published multiple sequence alignment (MSA) of 497 kinase domains (56). Ensembl BioMart (57), a web-based tool allowing the extraction of various biological data, was employed to match the kinase gene names to their associated Ensembl protein IDs. Furthermore, using BioMart, the PDB IDs of protein structures pertaining to the aggregated kinase protein IDs were retrieved, and the structural models matching these IDs were downloaded via the PDB’s bulk download tool in the CIF format. The workflow for download and classification of kinase structures and models is visible in Supp. Figure 1.

We predicted <u>AlphaFold2</u>models of kinase domains at a range of MSA depths. These models were generated using a modified implementation of the AlphaFold2_advanced (44) ColabFold repository. The specific modifications were that the ‘max_recycles’ and ‘is_training’ variables were set to 1 to reduce structural refinement and ‘True’ to enable model dropout, respectively. For modeling at each MSA depth, the max_msa variable was set to “{desired MSA depth}:{double desired MSA Depth}.” The maximum number of models output per kinase at each MSA depth was set to five. The resultant models were ranked by the pLDDT score calculated by AF2.

Memory limitations forbade successful predictions of models of MASTL at a depth of 512. All predicted models were visualized using the PyMOL Molecular Graphics System (version 2.5.4) hosted by Schrödinger, LLC.

### DFG-Out-Only Model Alignment-Based Prediction

A total of 161 sequences of kinases that were successfully modeled by AF2 in a DFG-out state at an MSA depth of 8 were extracted and combined in a new, custom MSA. This MSA was used in an attempt to predict structures of all kinase domains that were more abundant in DFG-out conformations. Variations in depths of this alignment (161, 32, 16, 8 and 4), using a random seed of 1, and randomly sampled kinase sequences from the original alignment of 161 sequences, were used to make new structure predictions for all kinases according to the same protocol described in the previous subsection.

### ESMFold Model Generation

Models were predicted based on sequences from the same multiple sequence alignment of 497 kinase domains (56) using the v1 implementation of ESMFold (downloaded from https://huggingface.co/facebook/esmfold_v1/tree/main) (2) with default parameters.

### Kinase Conformation Classification

Initial conformational classification of PDB structures and AF2 and ESMFold models of kinases were conducted using *Kincore* (42) and *Kinformation* (51). Due to its improved coverage of the kinome, *Kincore*’s results were shown and analyzed throughout this paper. This method classifies kinases into eight conformations based on the DFG motif and ɑC-Helix movements, which we grouped into the following six conformations to simplify classification by DFG-in and parse by ɑC-Helix conformation: DFG-in/ɑC-Helix-in (CIDI), DFG-in/ɑC-Helix-out (CODI) (DFG-in clusters in *Kincore*: BLAminus, BLAplus, ABAminus, BLBminus, BLBplus, and BLBtrans in *Kincore*), DFG-out/ɑC-Helix-in (CIDO), DFG-out/ɑC-Helix-out (CODO) (DFG-out cluster: BBAminus in *Kincore*), DFGinter (DFGinter in *Kincore*) and Unassigned (Unassigned in *Kincore*) (26). Kincore could not identify the residues of and near the DFG motif (or substituted residues) used for conformation classification for a total of twelve kinases, so these were excluded from the analysis. A total of 5,136 structures and 494 models were classified from Protein Data Bank and the AF2 Database, respectively.

Certain UniProt IDs were associated with multiple kinase models downloaded from the AF2 Database, rarely in more than one conformation; to calculate the fractional contribution to each conformation by each kinase containing these domains, 1,000 random samples of one conformation per kinase domain were taken from all conformations for each ID. These fractional contributions were calculated 10 times and averaged to calculate an overall contribution that was added to the corresponding conformation fraction in distribution calculations.

For simplicity, the most frequently observed conformation type was considered for family-wise distribution calculations. 2,425 AlphaFold2-predicted models were classified for each MSA depth, except for the depth of 512, for which 2,420 models were classified (due to memory limitations, AF2 failed to generate models for MASTL). A total of 486 ESMFold models were classified into the six conformations. Mapping of conformation counts to kinases and their groups were performed using KinMap (45). Comparison of kinase conformations represented in the PDB and at an MSA depth of 8, displayed in these maps, was performed by performed via a one-sided, paired t-test (p-value (Average_AF2_ > Average_PDB_)).

### Dihedral Plots

Plots of pseudo-dihedral angles used to visualize movement of the DFG motif were generated using Möbitz (49). Dihedral angles were calculated using the Cɑ carbons of 4 consecutive residues near or including the DFG motif. Biopython 1.81 (58) was used to calculate the coordinates of the required Cɑ carbons. The DFG Asp through the first residue after the DFG motif was used to define the dihedral on the X-axis, and the two residues preceding the DFG motif through the DFG Phe were used to define the same on the Y-axis.

### Statistical Analyses

Comparisons of fractions of specific conformations were performed by bootstrapping from distributions of data and counting the number of the conformation sampled, all 1,000 times. P-values of these comparisons were calculated using one-sided Wilcoxon rank-sum tests. The Chi-squared, Wilcoxon rank-sum, Fisher’s exact and paired t-tests used in our analyses were performed using the ‘chisq.test,’ ‘wilcox.test,’ ‘fisher.test,’ and ‘t.test,’ functions, respectively, from R 4.9.0. The specific analyses performed between different datasets are described in Supp. Figure 1.

### TM-Score Calculations

TM-Score is a metric used to compare structural similarity and is sensitive to smaller, local variations affecting the score of the global fold (46). TM-Scoring of AF2 models against PDB structures was performed using the ‘tmtools’ module and a series of helper functions obtained from https://gist.github.com/NicholasClark/. To simplify the analysis and focus on kinases with catalytic activity, TM-Score comparisons were only conducted between structures sharing a specific ‘DFG’ motif. Distributions of TM-Scores were calculated in terms of the averages of the highest TM-Scores for AF2 models compared to PDB structures of the same kinases for each MSA depth and structural conformation.

## Acknowledgments

This work was supported by the U01CA271318 grant from the National Cancer Institute. It was also supported in part through the computational and data resources and staff expertise provided by Scientific Computing and Data at the Icahn School of Medicine at Mount Sinai, as well as the Clinical and Translational Science Awards (CTSA) grant UL1TR004419 from the National Center for Advancing Translational Sciences. The authors thank Nicole Zatorski for assistance with various Python and R packages, as well as John Sekar for advice on statistical analyses.

