## Supplementary Figures and Tables for "Exploring the Druggable Conformational Space of Protein Kinases Using AI-Generated Structures"

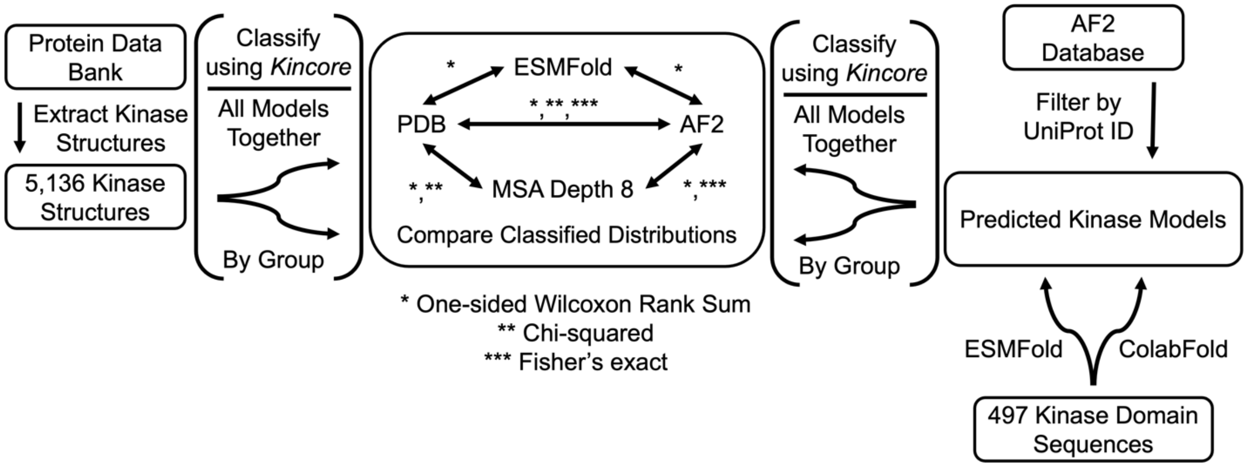


**Supplementary Figure 1. Workflow diagram for analysis of kinase structural models.** Kinase structures are downloaded from the Protein Data Bank (PDB) before being classified together and by their respective groups. The same is performed on kinase models downloaded from the AlphaFold2 Protein Structure Database ('AF2 Database') and generated through ColabFold and ESMFold. Distributions of structures and/or models are then compared via various statistical tests.

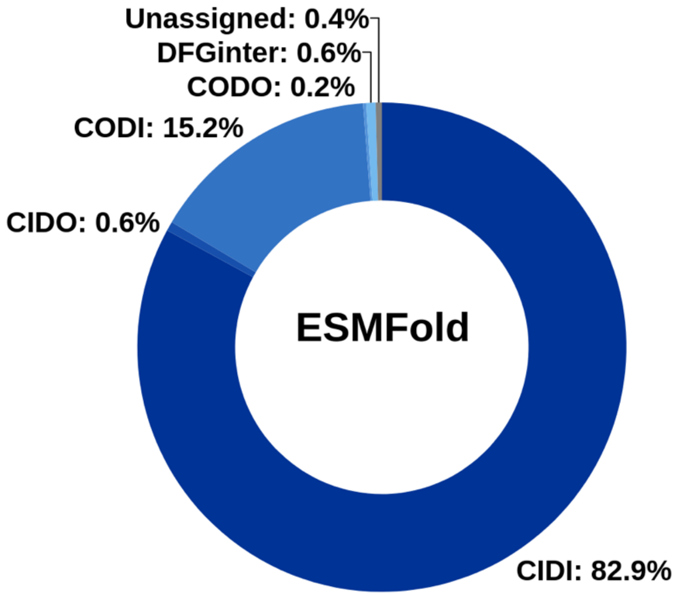


**Supplementary Figure 2. Distribution of human kinase models predicted by ESMFold by conformation (n = 486)**. Models generated by ESMFold and classified to a conformation by Kincore (42), as described in Methods. showed a strong preference for the active state (CIDI), significantly more so than the PDB (p-value $PDB_{CIDI}$ < $ESMFold_{CIDI}$ < $2.2 \times 10^{-16}$ or AlphaFold2 (p-value $AF2_{CIDI}$ < $ESMFold_{CIDI}$ < $2.2 \times 10^{-16}$), and similarly low preference for DFG-out states (CIDO and CODO), even lower than the PDB (p-value $PDB_{DFG\text{-}out}$ > $ESMFold_{DFG\text{-}out}$ < $2.2 \times 10^{-16}$) or AlphaFold2 (p-value $AF2_{DFG\text{-}out}$ > $ESMFold_{DFG\text{-}out}$ = $5.47 \times 10^{-281}$). P-values were obtained by a one-sided Wilcoxon rank-sum test.

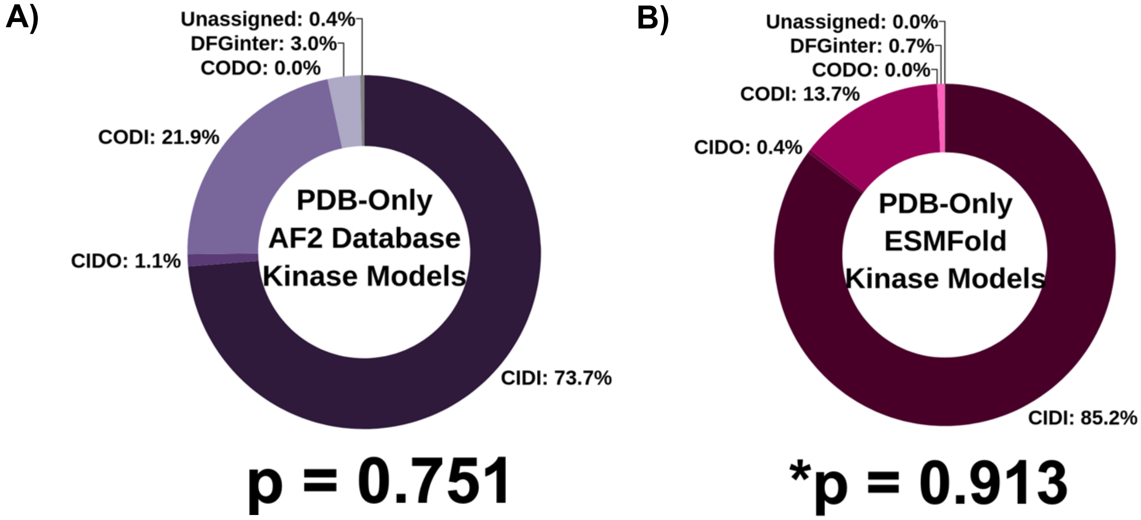


**Supplementary Figure 3. Distributions of conformations of only A) AF2 database and B) ESMFold models of human kinases with deposited experimental structures in the PDB.** The AF2 and ESMFold distributions were compared via a Fisher exact test. The resultant p-values of 0.751 and 0.913 for the AF2 Database and ESMFold overlap indicated that these distributions are not statistically different from all kinase models taken from the whole AF2 Database and ESMFold dataset, respectively.

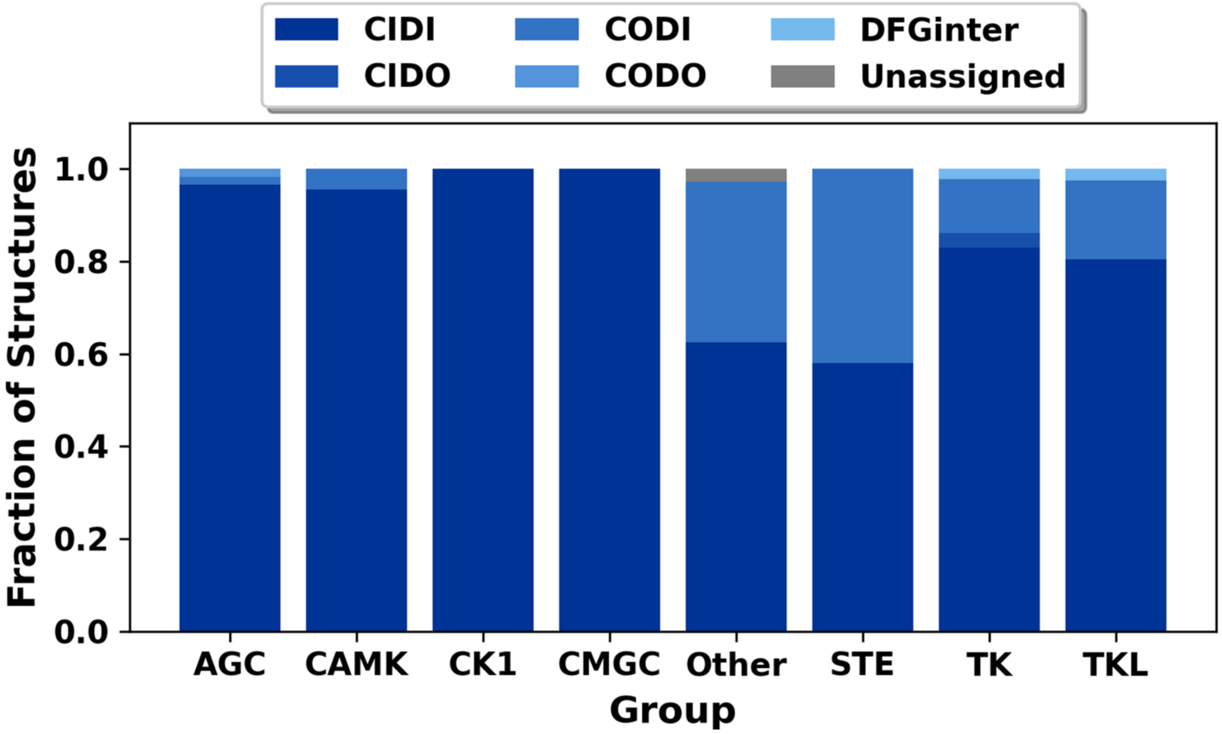


**Supplementary Figure 4. Fractional distributions of kinase conformations predicted by ESMFold by group.** Kinases are grouped into the following groups: AGC (PKA, PKG, PKC families;), CAMK (Calcium/calmodulin-dependent), CK1 (Casein kinase 1), CMGC (CDK, MAPK, GSK3, CLK families), STE (Sterile 7, Sterile 11, Sterile 20 kinases), TK (Tyrosine kinase; Tyrosine kinase-like), and Other. As in the PDB (Figure 1D) and AF2 Databases (Figure 1E), the active (CIDI) conformations were the most abundant across all families, while DFG-out conformations (i.e., CIDO, CODO) were substantially under-represented.

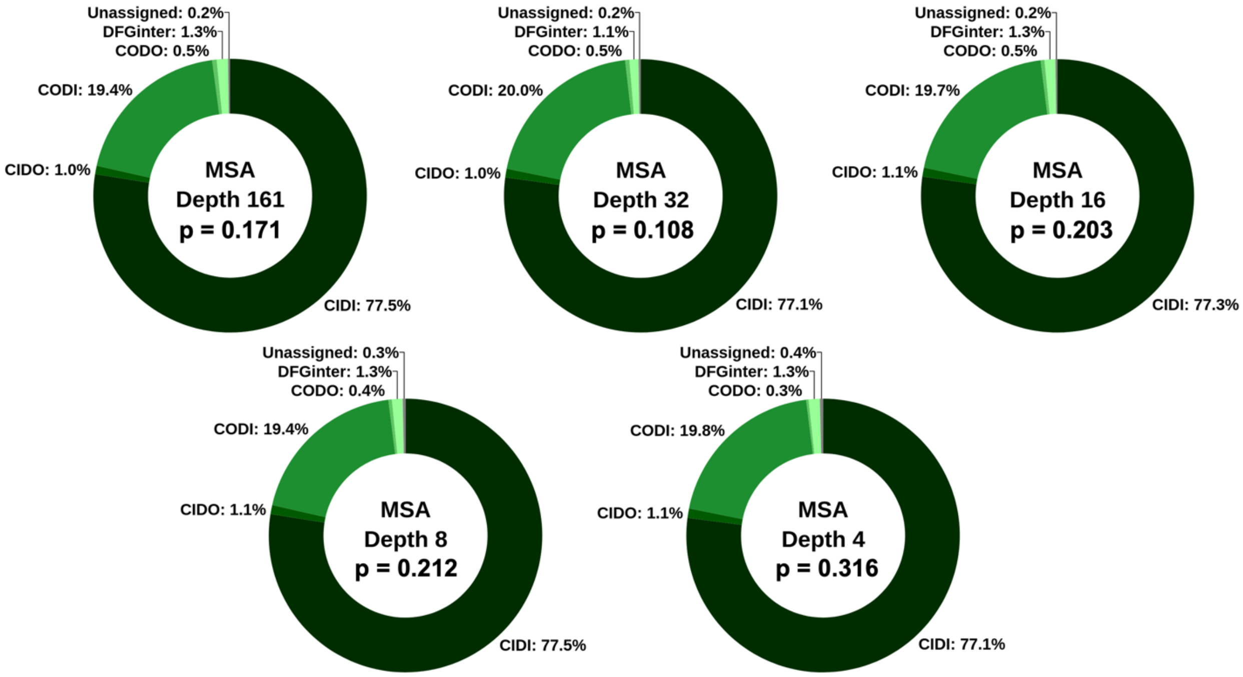


**Supplementary Figure 5. Distributions of conformations of AF2 models generated using a custom MSA.** A custom MSA was constructed from sequences of kinases predicted in at least one DFG-out conformation using AF2 at an MSA depth of 8. The counts of sequences in the custom MSA were gradually reduced to create new MSAs of various depths: 161, 32, 16, 8, and 4. Fisher's exact tests were used to compare each distribution to that of the AF2 Database, where all p-values indicated that none of these distributions were significantly different from those of models predicted using AF2's default parameters at a significance level of $p < 0.05$.

**Supplementary Table 1. Comparison of distributions of models predicted by AlphaFold2 at various MSA depths to that of the AlphaFold2 Database.** P-values were obtained using the Fisher's exact test.

| MSA Depth[a] | p-value[b] |
|---|---|
| 512 | 0.499 |
| 128 | 0.0045 |
| 32 | 0.0045 |
| 16 | 0.0045 |
| 8 | 0.0045 |
| 4 | 0.0045 |
| 2 | 0.0045 |

[a] MSA Depth corresponds to the number of sequences included in the Multiple Sequence Alignment (MSA) used as input for AF2 through ColabFold.

[a] At a significance level of 0.05, all p-values indicate that only the distribution at an MSA depth of 512 was statistically similar to that of the AF2 Database. All others were significantly different.

**Supplementary Table 2. Comparison of conformational fraction distributions of kinase models generated by AF2 at different MSA depths. P-values were calculated using the Chi-squared test.**

| MSA Depth[a] | 512 | 128 | 32 | 16 | 8 | 4 | 2 |
|---|---|---|---|---|---|---|---|
| 512 | | | | | | | |
| 128 | 1.73E-03 | | | | | | |
| 32 | 6.89E-21 | 6.59E-07 | | | | | |
| 16 | 2.76E-17 | 2.72E-11 | 1.02E-14 | | | | |
| 8 | 1.60E-209 | 1.79E-224 | 7.16E-253 | 5.72E-178 | | | |
| 4 | < 2.2E-16 | < 2.2E-16 | < 2.2E-16 | < 2.2E-16 | 4.72E-198 | | |
| 2 | < 2.2E-16 | < 2.2E-16 | < 2.2E-16 | < 2.2E-16 | < 2.2E-16 | 3.77E-122 | |

At a Bonferroni-corrected significance level of $p < 0.00238$, all distributions by each MSA depth are statistically different from each other.

[a] MSA Depth corresponds to the number of sequences included in the Multiple Sequence Alignment (MSA) used as input for AF2 through ColabFold.

**Supplementary Table 3. Comparison of conformation fractions of kinase models deposited in AF2 Database and predicted by AF2 at MSA depth 8.** P-values were calculated using the one-sided Wilcoxon rank-sum test at a significance level of 0.05.

| Group[a] | CIDI[b] | CIDO | CODI | CODO | DFGinter | Unassigned |
|---|---|---|---|---|---|---|
| AGC | < 2.2e-16 | < 2.2e-16 | 1 | 1 | < 2.2e-16 | < 2.2e-16 |
| CAMK | < 2.2e-16 | < 2.2e-16 | 1 | < 2.2e-16 | < 2.2e-16 | < 2.2e-16 |
| CK1 | < 2.2e-16 | < 2.2e-16 | < 2.2e-16 | < 2.2e-16 | < 2.2e-16 | < 2.2e-16 |
| CMGC | < 2.2e-16 | < 2.2e-16 | 1 | < 2.2e-16 | < 2.2e-16 | < 2.2e-16 |
| Other | < 2.2e-16 | 1 | 1 | < 2.2e-16 | < 2.2e-16 | < 2.2e-16 |
| STE | < 2.2e-16 | < 2.2e-16 | 1 | < 2.2e-16 | < 2.2e-16 | < 2.2e-16 |
| TK | < 2.2e-16 | 1 | 1 | < 2.2e-16 | 1 | < 2.2e-16 |
| TKL | < 2.2e-16 | 1 | 1 | < 2.2e-16 | < 2.2e-16 | < 2.2e-16 |

[a] Group indicates kinase group within which comparisons are being made.

[b] 'CIDI' indicates the significance of the $p(AF2_{CIDI} > 8MSA_{CIDI})$ comparison of 'CIDI' fractions. All other comparisons indicate comparison by $p(AF2_{CONF} < 8MSA_{CONF})$.
